# A protocol to automatically calculate homo-oligomeric protein structures through the integration of evolutionary constraints and ambiguous contacts derived from solid- or solution-state NMR

**DOI:** 10.1101/714857

**Authors:** Davide Sala, Linda Cerofolini, Marco Fragai, Andrea Giachetti, Claudio Luchinat, Antonio Rosato

**Affiliations:** Magnetic Resonance Center (CERM), University of Florence, Via Luigi Sacconi 6, 50019 Sesto Fiorentino, Italy; Consorzio Interuniversitario di Risonanze Magnetiche di Metallo Proteine, Via Luigi Sacconi 6, 50019 Sesto Fiorentino, Italy; Department of Chemistry, University of Florence, Via della Lastruccia 3, 50019 Sesto Fiorentino, Italy

## Abstract

Protein assemblies are involved in many important biological processes. Solid-state NMR (SSNMR) spectroscopy is a technique suitable for the structural characterization of samples with high molecular weight and thus can be applied to such assemblies. A significant bottleneck in terms of both effort and time required is the manual identification of unambiguous intermolecular contacts. This is particularly challenging for homo-oligomeric complexes, where simple uniform labeling may not be effective. We tackled this challenge by exploiting coevolution analysis to extract information on homo-oligomeric interfaces from NMR-derived ambiguous contacts. After removing the evolutionary couplings (ECs) that are already satisfied by the 3D structure of the monomer, the predicted ECs are matched with the automatically generated list of experimental contacts. This approach provides a selection of potential interface residues that is used directly in monomer-monomer docking calculations. We validated the protocol on tetrameric L-asparaginase II and dimeric Sod1.

## INTRODUCTION

Many proteins carry out their functional role acting as part of protein assemblies, i.e. a combination of different proteins (hetero-complexes) or of multiple copies of the same monomeric unit (homo-complexes). The assembly of the correct biological complex strongly depends upon specific protein-protein interactions (PPIs) that often are conserved among species (Qian et al., 2011; Sun and Kim, 2011). Frequently, an initial step in the study of an assembly is to characterize the three-dimensional structure of its individual subunit components either by X-ray crystallography or NMR spectroscopy. Among NMR techniques, solid-state NMR (SSNMR) has been receiving increasing attention because it is not limited by protein size, solubility, crystallization problems, presence of inorganic/organic matrices or lack of long-range order that often make the application of other structural biology methods unsuitable. In particular, it is straightforward to extend SSNMR experiments designed for individual proteins to the investigation of protein assemblies (Demers et al., 2018), as the quality of SSNMR spectra does not decrease with increasing molecular weight, as happens for solution NMR.

A crucial step in the application of SSNMR to structure determination is the identification and assignment of through-space nucleus-nucleus interactions. DARR (Dipolar Assisted Rotational Resonance) is a commonly used pulse sequence for this purpose, which is based on ^13^C-^13^C magnetization transfer through proton-driven spin diffusion (Takegoshi et al., 2001). Tuning of experimental DARR parameters allows users to select the range of distances at which inter-nuclear interactions are sampled. Although solid-state resonance lines of protein complexes are narrow, spectral congestion makes the assignment of DARR peaks a challenging task. In practice, DARR experiments yield a list of ambiguous contacts in which the quaternary contacts must be separated from intra-monomeric contacts to determine the 3D structure of the complex. In hetero-complexes this problem can be alleviated by using different schemes for enrichment in stable NMR-active isotopes (^13^C, ^15^N) in the various subunits of the complex (Göbl et al., 2014); for instance, one subunit can be uniformly enriched while all other subunits are not. This approach may not be very effective for homo-complexes, and more complex and labor intensive strategies for the asymmetric enrichment of all subunits have been proposed (Traaseth et al., 2008). Thus, the investigation of homo-complexes by SSNMR often remains a manual task, especially with respect to the identification of inter-subunit contacts.

Coevolution analysis assumes that evolutive pressure favors the preservation of protein function through the conservation of fundamental residue interactions (Salinas and Ranganathan, 2018). This concept has been implemented, among others, in global coevolutionary or direct coupling analysis (DCA) methods (Morcos et al., 2011; Weigt et al., 2008). These methods differ for the types of approximation used, from dimensional reduction (Cocco et al., 2013) to pseudo-likelihood maximization (Ekeberg et al., 2013) and others (Burger and van Nimwegen, 2010; Jones et al., 2012; Skwark and Elofsson, 2013). The information derived allows the identification of multiple protein conformational states (Morcos et al., 2013; Sutto et al., 2015) and the prediction of tertiary protein structures, either alone or in combination with experimental data (Anishchenko et al., 2017; Dago et al., 2012; Marks et al., 2012, 2011; Tang et al., 2015). Coevolution analysis can detect also ECs corresponding to inter-subunit contacts (Hopf et al., 2014; Ovchinnikov et al., 2014; Rodriguez-Rivas et al., 2016; Schug et al., 2009; Szurmant and Weigt, 2018). The identification of ECs consistent with PPIs for hetero-complexes requires the creation of a *joint* multiple sequence alignment (MSA) in which each line corresponds to an interacting protein pair (Bitbol et al., 2016; Burger and van Nimwegen, 2008; Cheng et al., 2014; Procaccini et al., 2011). This is a relatively complex task, especially due to the analysis required for the separation of orthologs and paralogs, prior to the construction of the MSA. Instead, the coevolution analysis of homo-complexes is based on a single protein sequence and thus on a single MSA. While this simplifies the construction of the alignment, it makes the identification of ECs belonging to inter-molecular contacts much more complicated because such information is hidden among hundreds or thousands of ECs of which the majority are tertiary contacts (dos Santos et al., 2015; Uguzzoni et al., 2017). The removal of tertiary contacts requires knowledge of the 3D structure of the monomeric protein. Notably, there is a relevant number (about 2000) of protein families annotated as forming homo-oligomeric assemblies *in vivo* with a deposited monomeric structure in the Protein Data Bank (PDB) (El-Gebali et al., 2019; Rose et al., 2015). These families potentially constitute an interesting target for homo-oligomeric structural predictions, also in the frame of drug discovery (Bai et al., 2016).

In the present work we developed a protocol to extract information on the protein-protein interface of homo-complexes from SSNMR-derived ambiguous contact lists, which can be automatically generated, using coevolution analysis. All the ECs with a relevant probability to be true residue interactions in either the monomer (intra-monomeric contacts) or in the homo-oligomerization interface (inter-monomeric contacts) are considered. The removal of intra-monomeric ECs requires the availability of the structure of the monomer. The predicted ECs with possible matches to experimental peaks are used to identify candidate interface residues. The final list of such residues is used directly in protein-protein docking calculations. The same protocol can be also applied using only solution-state NMR data.

## RESULTS

Our protocol aims to predict the structure of homo-oligomeric complexes by using ambiguous NMR contacts to identify reliable inter-monomeric contacts within the list of ECs. The whole procedure, which is described in detail in the next section, can be divided in two main parts. First, intra-monomeric evolutionary couplings (ECs) are removed from the list of ECs based on the 3D structure of the monomer. Second, the list of ECs predicted to potentially be at the complex interface is compared with the list of ambiguous NMR contacts to extract all residue pairs matching both the predicted and the experimental dataset. The protocol was validated by predicting the tetrameric structure of *Escherichia coli* L-asparaginase II (Cerofolini et al., 2019) (PDB ID: 6EOK), in which two distinct dimeric conformations must be recognized to reconstruct the functional complex (Fig. 1). Furthermore, the robustness of the procedure in the identification of complexes with small interface regions was tested by predicting the structure of dimeric human apo Sod1 (Bertini et al., 2009) (PDB ID: 3ECU) (Fig. 1). For L-asparaginase II we used solid-state NMR data (Cerofolini et al., 2019), whereas for Sod1 we used solution NMR data (Bertini et al., 2009).

**Figure 1.**
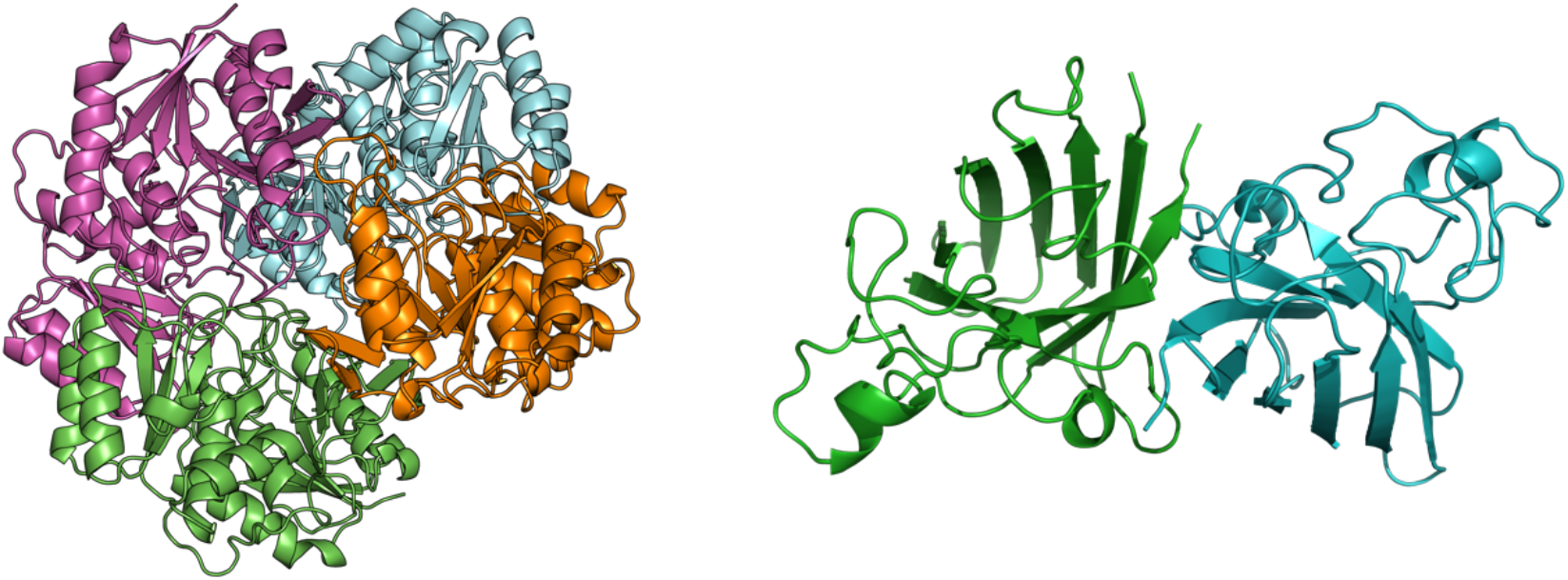
Crystal structures of the tetrameric L-asparaginase II and the dimeric apo Sod1.

### Description and application of the protocol

This protocol calculates a list of putative interface residues to be used as input to HADDOCK for docking calculations. It needs four inputs (Fig. 2): one or more files with the list of ECs, the structure of the monomer, the experimental NMR-derived list of ambiguous contacts and the Naccess file (rsa format) with the per-residue relative solvent accessible area. The ECs of the target protein are obtained from so-called coevolution analysis. A number of servers performing coevolution analysis are available online (see *Methods*). In general, they need the protein sequence as input to predict a contact map from multiple sequence alignments (MSAs). The output is a list of residue pairs scored for the probability that they are actually in contact in the monomeric or oligomeric structure. We apply a probability cutoff P to remove ECs with low probability of being true interactions. Coevolution analysis usually outputs from hundreds to thousands of ECs that cannot be assigned as intra-monomeric or inter-monomeric contacts without any structural information. As a consequence, our protocol calculates for each EC the corresponding Cα-Cα distance in the 3D structure of the monomer and all the ECs below the distance cutoff of 12 Å are classified as intra-monomeric and removed.

**Figure 2.**
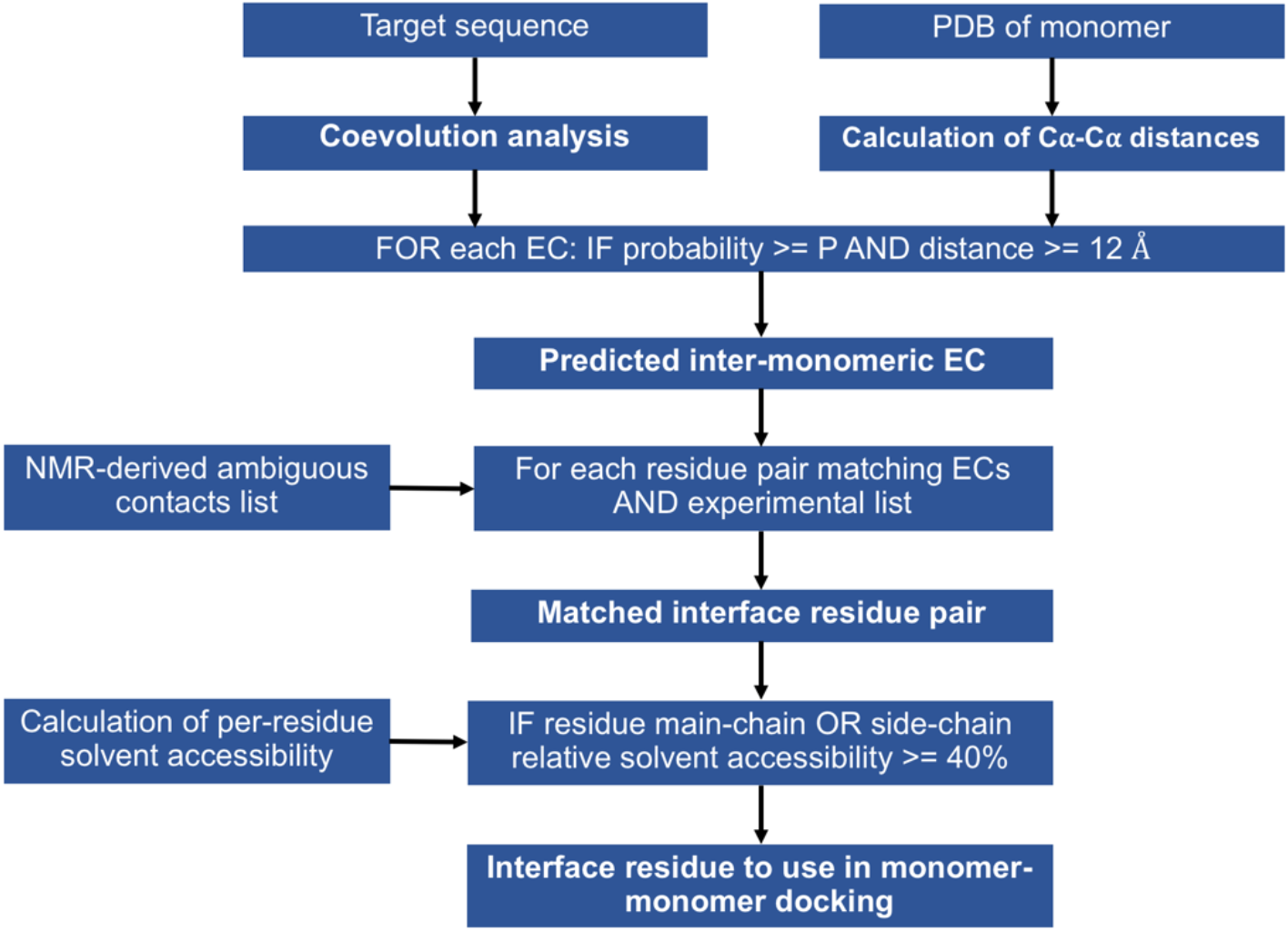
Scheme of the protocol adopted to predict the structure of homo-oligomeric complexes using coevolution analysis and ambiguous NMR contacts.

After the removal of intra-monomeric ECs, the resulting list is enriched in contacts across the interaction interface (inter-monomeric ECs). Nevertheless, it still contains a relevant number of false-positives. False-positives can be either ECs that do not correspond to a true residue-residue interaction or ECs that correspond to intra-monomeric interactions that occur in conformations sampled during the physiological conformational dynamics of the protein. The EC list thus cannot be used directly in docking calculations. We thought that the rate of false positives could be reduced by leveraging the information present in the list(s) of ambiguous contacts provided by NMR experiments. Indeed, NMR-derived contacts list of protein complexes are affected by a high level of ambiguity caused by the accidental overlap of NMR resonances, making the extraction of reliable inter-monomeric contacts an arduous task. Our protocol overcomes this bottleneck by matching the predicted inter-monomeric ECs with the experimental list to extract information present in both the datasets. In practice, residue pairs in the predicted inter-monomeric EC list are matched to ambiguous assignments in the experimental list, providing a list of interface residue pairs.

The number of residual false-positives in the matched list is further decreased by removing all the residues with a relative solvent accessibility lower than 40% in both main-chain and side-chain (i.e. buried residues). The remaining residues constituting the output list from our protocol can be used directly as ambiguous interaction restraints (AIRs) in monomer-monomer docking calculations with HADDOCK. The protocol can be run using the python script provided as supplementary material (*SI Appendix*).

We assessed the accuracy of the protocol in predicting residues at the homo-oligomeric interface for different probability cutoffs (Tables 1 and 2). Furthermore, we evaluated the NMR data contribution to the prediction accuracy by comparing the results obtained with or without (“ECs + NMR” and “ECs only”, respectively) matching with the NMR data. A residue accurately predicted at the complex interface is defined as a true-positive (TP) residue. More in detail, we defined a true-positive (TP) residue as having at least one atom with a distance < 7 Å from any atom located on a different chain in the crystal structure of the complex.

**Table 1.** Number of residues predicted to make contacts across the L-asparaginase II homomeric interface. The protocol was applied as depicted in figure 2 with the ECs matched with the NMR data “ECs + NMR” and without the matching step with NMR data “ECs only”. P indicates the probability threshold used to accept ECs. PPV = TP/(TP+FP).

| P | L-asparaginase II |  |  |  |  |  |
| --- | --- | --- | --- | --- | --- | --- |
|  | ECs only |  |  | ECs + NMR |  |  |
|  | TP+FP | TP | PPV | TP+FP | TP | PPV |
| **0.90** | 13 | 10 | 0.8 | 3 | 3 | 1.0 |
| **0.85** | 23 | 20 | 0.9 | 3 | 3 | 1.0 |
| **0.80** | 30 | 21 | 0.8 | 3 | 3 | 1.0 |
| **0.75** | 34 | 24 | 0.8 | 3 | 3 | 1.0 |
| **0.70** | 38 | 27 | 0.8 | 4 | 4 | 1.0 |
| **0.65** | 41 | 30 | 0.8 | 4 | 4 | 1.0 |
| **0.60** | 47 | 31 | 0.7 | 4 | 4 | 1.0 |
| **0.55** | 51 | 33 | 0.7 | 4 | 4 | 1.0 |
| **0.50** | 60 | 36 | 0.7 | 4 | 4 | 1.0 |
| **0.45** | 73 | 42 | 0.7 | 4 | 4 | 1.0 |
| **0.40** | 84 | 47 | 0.6 | 5 | 5 | 1.0 |
| **0.35** | 97 | 52 | 0.6 | 7 | 7 | 1.0 |
| **0.30** | 105 | 54 | 0.6 | 9 | 8 | 0.9 |
| **0.25** | 121 | 60 | 0.6 | 19 | 16 | 0.8 |
| **0.20** | 128 | 60 | 0.6 | 34 | 28 | 0.8 |

**Table 2.** Number of residues predicted to make contacts across the Sod1 homomeric interface.

| P | Sod1 |  |  |  |  |  |
| --- | --- | --- | --- | --- | --- | --- |
|  | ECs only |  |  | ECs + NMR |  |  |
|  | TP+FP | TP | PPV | TP+FP | TP | PPV |
| **0.90** | 0 | 0 | NA | 0 | 0 | NA |
| **0.85** | 0 | 0 | NA | 0 | 0 | NA |
| **0.80** | 0 | 0 | NA | 0 | 0 | NA |
| **0.75** | 4 | 3 | 0.7 | 0 | 0 | NA |
| **0.70** | 4 | 3 | 0.7 | 0 | 0 | NA |
| **0.65** | 4 | 3 | 0.7 | 0 | 0 | NA |
| **0.60** | 8 | 4 | 0.6 | 2 | 0 | NA |
| **0.55** | 10 | 4 | 0.4 | 2 | 0 | 0.0 |
| **0.50** | 17 | 5 | 0.2 | 4 | 0 | 0.0 |
| **0.45** | 23 | 7 | 0.3 | 5 | 1 | 0.2 |
| **0.40** | 29 | 9 | 0.3 | 5 | 1 | 0.2 |
| **0.35** | 38 | 12 | 0.3 | 9 | 3 | 0.3 |
| **0.30** | 50 | 14 | 0.3 | 18 | 7 | 0.4 |
| **0.25** | 68 | 17 | 0.2 | 27 | 7 | 0.3 |
| **0.20** | 74 | 17 | 0.2 | 48 | 9 | 0.2 |

In the case of the L-asparaginase II protein, the crystallographic complex is formed by four subunits with a D_2_ symmetry. Thus, the ensemble of all TP residues contains the amino acids at both dimeric interfaces. For this system, the inclusion of NMR data enhances the positive predictive value (PPV), defined as true-positive (TP) residue predictions over all predictions [TP/(TP+FP)], at all the probability cutoffs assessed (Table 1). In fact, on the basis of the “ECs only” analysis the absolute number of TP residues present in the prediction is significantly higher than the number of TP obtained after the match with NMR data. However, the same analysis also outputs a much greater number of FPs. Consequently, the “ECs + NMR” analysis features a PPV of 100% for P >= 0.35; the PPV remains very high (>= 80%) even at low probabilities (P < 0.35) and the number of predicted interface residues is sufficient to successfully drive docking calculations (see next section).

Instead, the Sod1 complex contains two subunits with a C_2_ symmetry and a small protein-protein interface. As a consequence, in the central part of the interface the inter-monomeric contacts involve residue pairs that also are at intra-monomer distance smaller than the 12 Å threshold that we used to remove intra-monomeric ECs. In practice, this structural organization significantly reduces the number of detectable TPs because the aforementioned inter-monomeric contacts are discarded. Furthermore, small interfaces are harder to predict computationally and also provide a lower number of NMR-detectable contacts. All these features make the Sod1 system challenging but useful to test the limits of the protocol. When considering the Sod1 protein, the “ECs only” protocol yielded a reasonable PPV for P >= 0.55, but with only a handful of TPs in the prediction (Table 2). Instead, the match with NMR data removed the signal for P>= 0.45 while retaining information at lower P values, especially for P = 0.30.

These results suggest that the quality of the initial EC prediction is quite important for the performance of our protocol, leading to a larger enhancement of the PPV when the prediction includes a larger number of TPs. When the EC data yielded is weaker and mixed with noise, our protocol retains a good part of the available information but the PPV is mostly unchanged.

### HADDOCK calculations for L-asparaginase II

The ECs at the P cutoff of 0.25 were matched with a solid state 2D^13^C-^13^C DARR dataset (mixing time 200 ms) holding 4937 ambiguous assignments, resulting in 19 surface residues predicted to be at the protein-protein interface (corresponding to 14% of the whole protein surface). The final 200 water-refined models generated by HADDOCK were analyzed by measuring the RMSD from the structure with the lowest HADDOCK score. The clustering algorithm grouped the models in 7 clusters (Fig. 3A). The first cluster was the most populated and included the models with the lowest score. Indeed, the lowest HADDOCK score model of the first cluster was a dimer with an RMSD of 0.7 Å from the crystallographic dimer formed by chain A and chain C of the tetrameric protein (Fig. 3B). In addition to the HADDOCK score, the desolvation energy calculated using empirical atomic solvation parameters proved to be an useful scoring function(Fernández-Recio et al., 2004), allowing the identification of the correct A-C dimer (Fig. S1).

**Figure 3.**
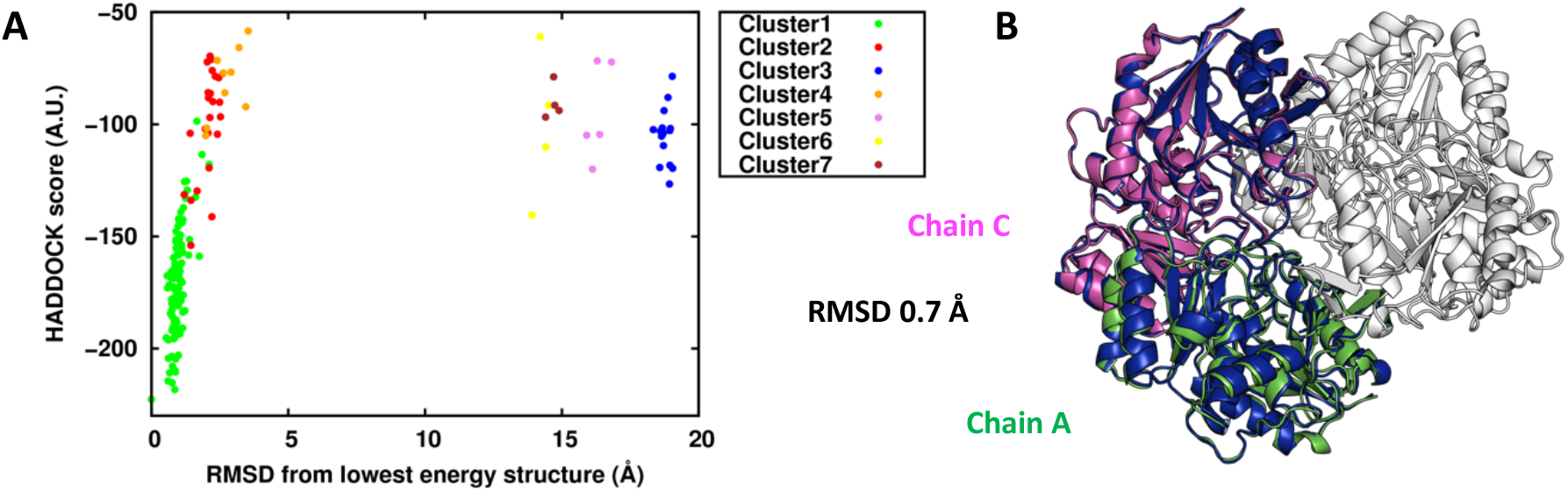
L-asparaginase II monomer-monomer docking. **A**) Plot of the HADDOCK score vs RMSD clusters distribution with respect to the lowest HADDOCK score model. **B**) Structural alignment between the lowest HADDOCK score model (in blue) of the first cluster and the crystal structure.

Both the predicted inter-monomeric ECs and the experimental NMR inter-monomeric contacts include residue pairs belonging to all the pairs of chains effectively in contact in the functional complex. In the case of the tetrameric L-asparaginase II, besides the largest A-C interface also chains A and D share a relevant number of contacts. According to this, in a single docking run one might expect to sample both relevant dimeric configurations (A-C and A-D) in two different clusters. Indeed, by checking the position of the 19 predicted interface residues within the crystal structure, it appears that the A-C and A-D interfaces were both mapped (Fig. 4). In fact, the largest portion of residues effectively in contact belonged to dimer A-C and the smallest portion to dimer A-D.

**Figure 4.**
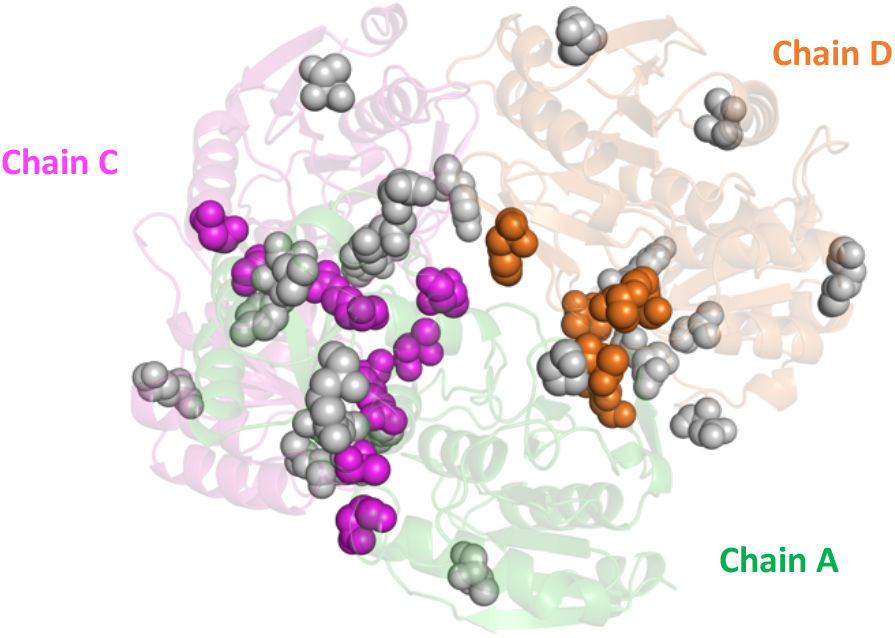
Projection on the crystal structure of the L-asparaginase II residues used to generate AIRs in the docking calculation. The residues making inter-monomeric contacts are shown as colored spheres (A-C interface in purple; A-D interface in orange).

However, the structural configuration present in the other clusters did not correspond to the A-D dimer. This could be easily verified by observing that the superimposition of the two dimers on the common chain A resulted in evident steric clashes between the subunits, as shown for the cluster 3 (Fig. 5). If the two dimers actually corresponded to the A-C and A-D dimers of the tetrameric structure, the superimposition on the A chain would have caused no significant clashes.

**Figure 5.**
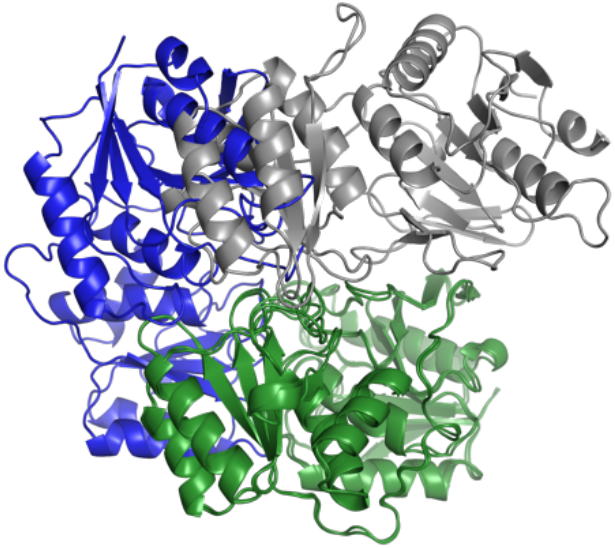
Superimposition on chain A (in green) of the third (in gray) and the best (in blue) dimer configurations in the first run.

In principle, the absence of the second compatible dimer in calculations can be due to two reasons. First, the interface residues belonging to the second configuration were not present in the AIRs dataset. Second, the residues belonging to the second interface region were present, but the correct structural configuration had a HADDOCK score worse than the wrong sampled configurations. In the present case, the latter reason was the relevant one. In fact, the wrong dimer models in general contained some contacts from both interface regions, thus satisfying a higher number of AIRs than the correct dimer A-D.

To obtain a model of the A-D dimer, we performed a second docking run in which the restraints already satisfied in the best cluster (containing the most favored configuration) of the first run were removed from the input dataset. To this end, we looked at the violation analysis of HADDOCK, and retained all contacts that were not satisfied by the majority of the members of the first cluster by at least 3 Å. This resulted in 9 residues being used as input to a second monomer-monomer docking run. As in the previous calculation, the first cluster was the largest and contained the models with the best HADDOCK score and desolvation energy (Fig. 6A and S2). Superimposing the lowest HADDOCK score water-refined model with the crystal structure resulted in an RMSD of 0.9 Å from the dimer A-D (Fig. 6B).

**Figure 6.**
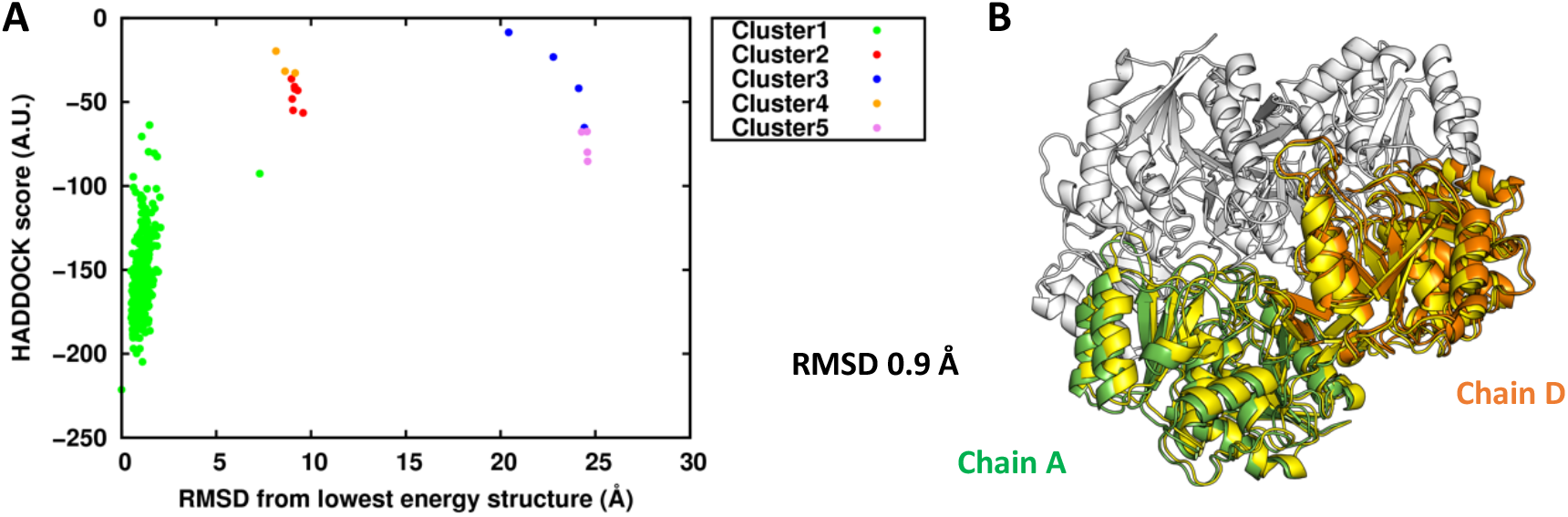
L-asparaginase II monomer-monomer docking using AIRs violated in the A-C dimeric model. **A**) Plot of the HADDOCK score vs RMSD clusters distribution with respect to the lowest HADDOCK score model **B**) Structural alignment between the lowest HADDOCK score model (in yellow) of the first cluster and the crystal structure.

In summary, the two correct dimeric conformations A-C and A-D were obtained performing two distinct docking runs, the first one with the whole AIRs dataset and the second one with the subset resulting from the removal of the AIRs satisfied in the best cluster of the first run. Crucially, this procedure provided us with two compatible non-overlapping dimeric models that, for symmetry, can be used to reconstruct the tetrameric model (Fig. 7). This step strictly depended by the correct identification of the structural model on which the distance violation analysis was carried out. In fact, selecting the third cluster of Fig. 3 to perform the violation analysis instead of the best one resulted in a second docking run that sampled again the dimer A-C in the two best clusters and not-compatible structural configurations in the others (Fig. S3).

**Figure 7.**
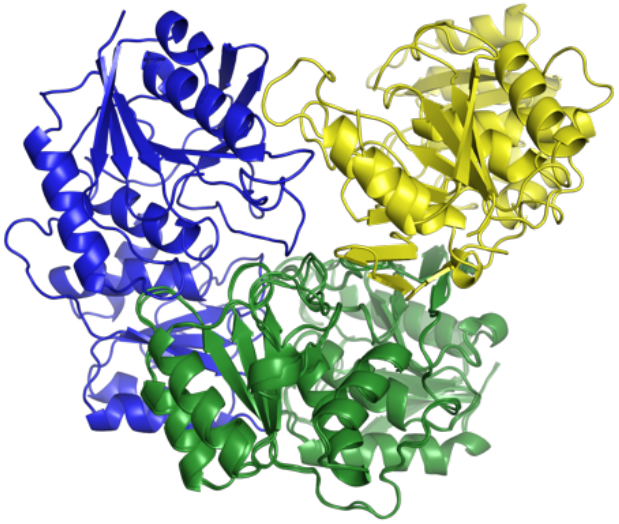
Superimposition on the chain A (in green) of the best structural configurations in the second run (in yellow) and in the first run (in blue).

Extracting the monomer from the PDB of the complex results in a protein model with the side chains oriented in a contact-ready state that favors the correct assembly, in terms of both docking score and RMSD from the experimental structure, as compared to incorrect docking poses. Thus, to test our protocol in a more realistic condition we generated 15 homology models of L-asparaginase II using the structure of the homolog from *Wolinella succinogenes*(Lubkowski et al., 1996) as the structural template (PDB ID 1WSA, chain A). The homology models had a backbone RMSD lower than 1 Å from the crystal structure of the *E. coli* protein, but widely differing in the orientation of the surface side chains. Each model was used in protein-protein docking with the same input AIRs of the “crystal P 0.25” runs, for both the A-C and A-D dimers. The results of Table 3 show the significant influence of the orientation of side chains on the ability of the docking calculations to sample the correct dimer in the best cluster. Based on the HADDOCK score of the best cluster for each model, the AC runs pointed out that the five runs with the best score also had the lowest RMSD from the crystal A-C dimer, (green gradient in the table). However, for these five models the second calculation with the AIRs providing the A-D dimer resulted in wrong dimeric conformations. Nevertheless, by inspecting the results for all models (Table 3), it turned out that the runs with the best HADDOCK scores (for their first clusters) indeed provided results conformations close to the crystallographic A-D dimer (in particular models 6 and 15). For further comparison, we performed a docking run of the crystallographic monomer with the 34 residues (25% of the whole protein surface) output by the protocol run at a P cutoff of 0.20. Changing the AIRs dataset with a larger one having the same PPV did not significantly affect the results.

**Table 3.**
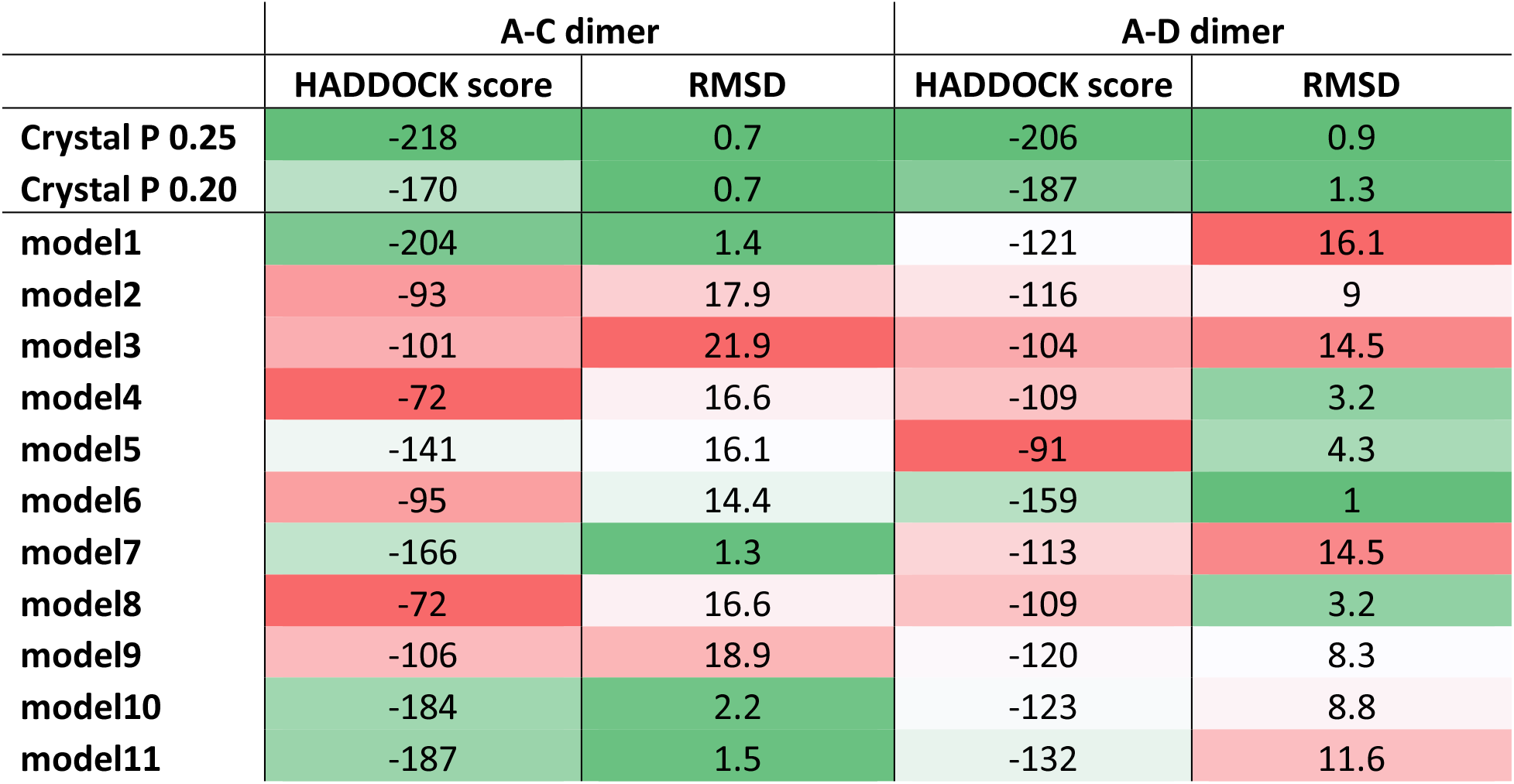
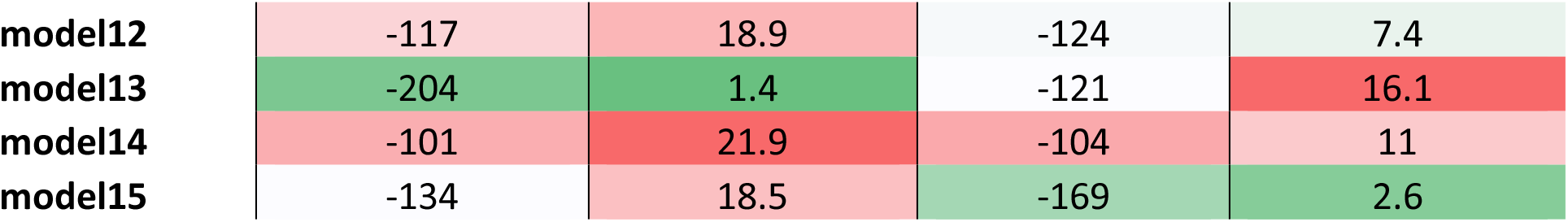
Docking results for homology models of L-asparaginase II. The two “Crystal” runs were performed using the chain A of the crystal structure. Each model mainly differs in the orientation of side chains. For each run the HADDOCK score of the best cluster (calculated as the average value of the 4 best structures of the cluster) and the RMSD of its best structure from the experimental dimer are reported.

|  | A-C dimer |  | A-D dimer |  |
| --- | --- | --- | --- | --- |
|  | HADDOCK score | RMSD | HADDOCK score | RMSD |
| **Crystal P 0.25** | -218 | 0.7 | -206 | 0.9 |
| **Crystal P 0.20** | -170 | 0.7 | -187 | 1.3 |
| **model1** | -204 | 1.4 | -121 | 16.1 |
| **model2** | -93 | 17.9 | -116 | 9 |
| **model3** | -101 | 21.9 | -104 | 14.5 |
| **model4** | -72 | 16.6 | -109 | 3.2 |
| **model5** | -141 | 16.1 | -91 | 4.3 |
| **model6** | -95 | 14.4 | -159 | 1 |
| **model7** | -166 | 1.3 | -113 | 14.5 |
| **model8** | -72 | 16.6 | -109 | 3.2 |
| **model9** | -106 | 18.9 | -120 | 8.3 |
| **model10** | -184 | 2.2 | -123 | 8.8 |
| **model11** | -187 | 1.5 | -132 | 11.6 |
| model12 | -117 | 18.9 | -124 | 7.4 |
| model13 | -204 | 1.4 | -121 | 16.1 |
| model14 | -101 | 21.9 | -104 | 11 |
| model15 | -134 | 18.5 | -169 | 2.6 |

Overall, the results described above pointed out the importance of generating a sufficiently large number of homology models to sample many different side chain orientations, thus increasing the probability to capture the orientation permitting residue-residue contacts across the monomeric interface. The best clusters of the two crystal runs showed that ideal side chain orientations provided the top HADDOCK score values. In line with this, the models that had the best HADDOCK scores resulted in the configurations closest to the crystal structure, with a backbone RMSD between 1 and 3 Å from it. For these models, the HADDOCK scores themselves were similar to the values observed for the runs starting from the crystal monomer. Indeed, superimposing on the chain A the AC dimer of model 13 and the AD dimer of model 15 or model6 showed two compatible dimeric models that, taken together, can be used to reconstruct the tetrameric structure (Figure S4)

### HADDOCK calculations for Sod1

The predicted inter-monomeric ECs at P=0.30 were matched with 7611 ambiguous assignments from solution-state 3D ^1^H ^15^N NOESY-HSQC spectrum. The protocol yielded 18 putative interface residues, corresponding to 23% of the whole monomer surface. By comparing the prediction to the of the crystal structure, it appeared that 7 out of 18 residues effectively formed inter-monomeric contacts (Fig. S5).

From the docking calculation starting with the crystal monomer we obtained 7 clusters with comparable HADDOCK score values (Fig. 8A). However, the distribution of the desolvation energies discriminated the second cluster as the most favored (Fig. 8B). Indeed, the structural alignment of the best model of this cluster with the experimental dimer revealed an impressive RMSD of 0.6 Å (Fig. S6A). Instead, the same superimposition on the crystal structure of the first cluster resulted in a dimer in which one of the two monomeric units was rotated by 180° with respect to the corresponding experimental monomer, while preserving the same interface region (Fig. S6B).

**Figure 8.**
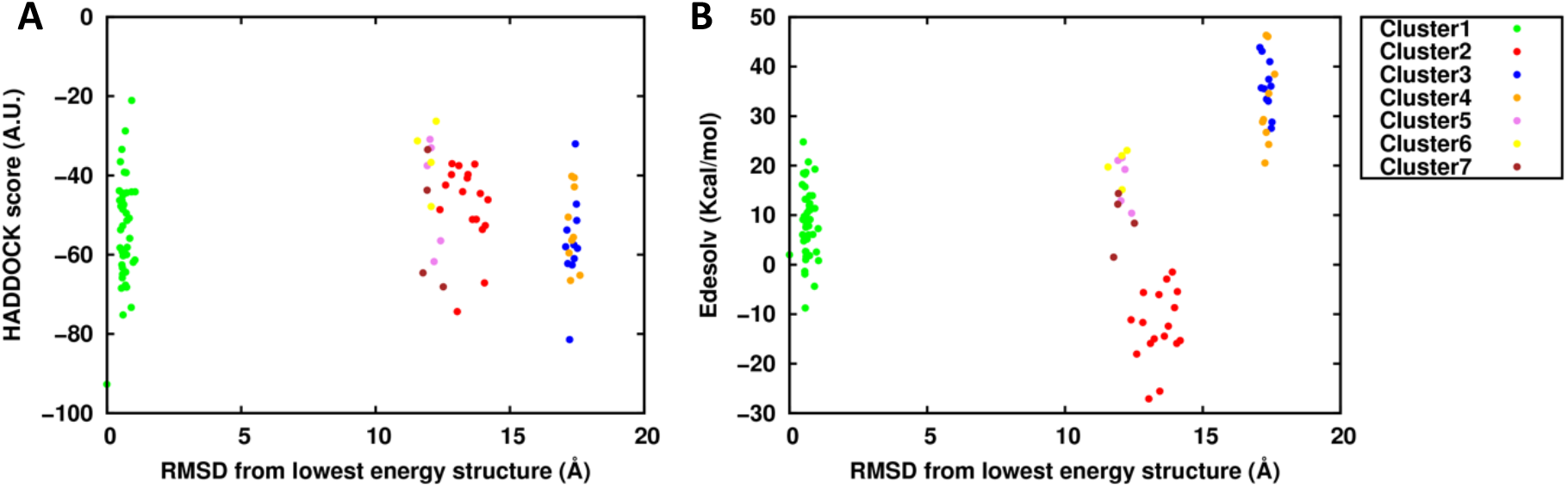
Sod1 clusters distribution with respect to the lowest HADDOCK score model. **A**) HADDOCK score distribution. **B**) Desolvation energy distribution.

## DISCUSSION

Solid State NMR is an attractive technique to study large protein assemblies as even systems with high molecular weight can provide very good spectra. However, the determination of their 3D structure involves two very time-consuming steps: the assignment of the side chains in contact at the interface between the subunits and, for homo-oligomeric complexes, the discrimination of intra- vs inter-monomer contacts. In particular, the correct identification of inter-monomer contacts usually requires extensive efforts by an experienced user. From the bioinformatics point of view, focusing on homo- rather than hetero-oligomers makes the interpretation of coevolution signals harder. In fact, the difficult step in the coevolution analysis of hetero-oligomers is the proper pairing of orthologs of interacting proteins and the corresponding removal of paralogs. Once this has been achieved, the creation of a *joint* MSA in which each line contains a pair of interacting proteins allows the straightforward use of predicted inter-protein contacts as restraints to drive the modelling of the quaternary structure (Bitbol et al., 2016; Hopf et al., 2014; Ovchinnikov et al., 2014). Instead, the coevolution analysis of homo-oligomers is based on a single protein MSA, which is relatively effortless to build. Unfortunately, the availability of the three-dimensional structure of the monomeric unit is necessary to successfully separate intra-monomeric and inter-monomeric ECs (Uguzzoni et al., 2017). In this work, we developed a protocol to integrate ECs with NMR-derived ambiguous contacts in order to identify interface residues in homo-oligomers. The input lists of ambiguous contacts can be automatically generated from appropriate solution or solid-state NMR spectra. Our protocol was validated by predicting two difficult cases: the tetrameric L-asparaginase II, in which two distinct dimeric conformations must be recognized to reconstruct the functional complex and the dimeric Sod1, in which the interface region is comparatively small.

The correct identification of interface residues was readily verified by comparing the output of the protocol with the known interfaces in the crystal structures of the two systems (Tables 1 and 2). This analysis showed that NMR data can be beneficial by enriching the predictions in true contacts (i.e. higher PPV). This improvement comes at the cost of reducing the absolute number of predicted residues, which however did not limit the subsequent docking calculations. The requisite for the integration of ECs and NMR data to be successful is that the initial list of potential inter-monomeric ECs contains enough information. This is clearly exemplified by the case of Sod1, for which the absolute number of predictions, after removing all contacts that could be satisfied within the monomer, was quite low. Consequently, many NMR signals could not be matched and the benefit in PPV was modest. Nevertheless, when the total number of predicted interface residues is in a reasonable range (15%-20% of all surface residues, i.e. 12-16 residues for Sod1) the prediction resulting from the integration of ECs and NMR data is more reliable than that based only on ECs.

To generate a 3D structural model of the oligomer, the output of our protocol can be exploited in docking calculations. As a proof-of-principle, we run these calculations starting from the monomer conformation observed in the crystal structure. This is an ideal case, where all the side chains at the protein-protein interface are already in the correct rotameric state to engage in the formation of the complex. All the same, it is important to perform this step to ensure that the output contains enough information to successfully drive the docking. This was indeed the case for the main dimer of L-asparaginase II (A-C) as well as for Sod1. The calculation with the complete AIR dataset could not identify the A-D dimer even though the dataset contained contacts belonging to both interfaces. The A-D interface is somewhat smaller than the A-C interface; as HADDOCK aims to satisfy the highest number of AIRs, the situation where the second chain of the dimer is positioned in between the two interfaces, thus partly satisfying both subsets of AIRs, is favored over the situation in which all of the A-D and none of the A-C AIRs are satisfied. To circumvent this bottleneck, it is necessary to separate the residues belonging to each interface. This was done by removing the contacts already satisfied in the first docking calculation to run a second calculation only with the unsatisfied restraints. The best cluster of the second run indeed matched closely the A-D dimer of the tetramer (Fig 6). Intriguingly, the AIRs derived from ECs only at a P cutoff of 0.8 (Table 1), whose number was similar to the number of AIRs used in the “ECs + NMR” calculations, did not contain information on the A-D dimer interface (not shown). Thus, the information provided by ECs at high levels of confidence is not balanced over the two interfaces, presumably due to the evolutionary history of the system. This makes it necessary to use data at lower P cutoffs, which is efficiently filtered by the ambiguous contacts provided by solid state NMR. The experimental data in fact contain information on both interfaces and thus is useful to extract both sets of true contacts from the list of ECs.

In a more realistic scenario one would use a homology model of the monomer as the input structure to docking calculations. We tested this scenario by generating 15 different models of L-asparaginase II (Table 3) and using the same input AIRs used in the docking of the crystal monomer for all calculations, so that the structure was the only source of variability. For the A-C dimer, we observed that in four cases the best model of the adduct was within 2 Å from the crystal structure, while an additional calculation provided a model with a RMSD of 2.2 Å. The A-D dimer resulted in a similar situation, with two structures within 3 Å and another two at 3.2 Å. Remarkably, there was a very good correlation between the HADDOCK score and the RMSD, allowing the more accurate models to be identified quite straightforwardly. It is also noteworthy that the best results obtained with the homology models had scores close to those obtained with the crystal monomer, which can be reasonably assumed to represent the best possible score. It thus appears that sampling a relatively extensive ensemble of different conformations is an important factor to obtain accurate models of the oligomer in a real-life setting.

In summary, our protocol allowed us to predict homo-oligomeric structure in multimers and in presence of a small homodimerization interface. Notably, this goal was achieved with a minimal user effort, making the determination of the 3D structure of the complex faster than using experimental data alone. The only parameter that must be decided by the user is the probability cutoff P below which the ECs are removed. In our hands selecting a P cutoff such that the number of predicted interface residues was 15%-20% of the number of surface residues in the monomer worked well. The results of our protocol clearly depend upon the quality of the ECs obtained from the online servers. Their integration with NMR data serves two different purposes, namely enriching the input AIRs in true contacts when working at low P cutoffs and removing biases among different regions of the protein. From the point of view of NMR spectroscopists, the present work provides a methodology to analyze homo-oligomers with reduced manual effort.

## METHODS

### Computational aspects

The protocol described in the “results” section can be carried out running the python script provided (*SI Appendix*). The script needs four inputs: the ECs files, the PDB structure of the monomeric protein, the experimental ambiguous NMR contacts list and the Naccess file (rsa format) with the relative solvent accessibility of the residues. Details about inputs preparation, script steps, and docking protocol adopted for the L-asparaginase II and Sod 1 are described below.

The ECs for both proteins were collected using 3 servers available online: Gremlin (Ovchinnikov et al., 2014) (http://gremlin.bakerlab.org), RaptorX (Wang et al., 2017; Xu et al., 2016) (http://raptorx.uchicago.edu/) and ResTriplet (Yang Li, Chengxin Zhang, Dongjun Yu, 2018) (https://zhanglab.ccmb.med.umich.edu/ResTriplet/). The last two methods are supervised but the PDBs used in this work were not present in the training sets. The MSA in the Gremlin server was generated with the Jackhmmer method and default options (Eddy, 1998). Using different servers adopting different methods in the ECs generation can result in multiple copies of the same EC with different computed likelihood probability. If this is the case, the EC with the highest probability is kept.

The reference protein structures were retrieved from the Protein Data Bank: *E. coli* L-asparaginase II corresponds to PDB ID 6EOK, whereas human apo-Sod1 has the PDB ID 3ECU. Inter-monomeric ECs were identified by removing from the full EC lists all residue pairs with a corresponding Cα-Cα distance < 12 Å in chain A of the structures. This distance was already proved as an excellent threshold in the selection of true contacts across the interface (Uguzzoni et al., 2017).

The experimental procedure for the generation of the ambiguous NMR contacts list is described in the next section.

The per-residue relative solvent accessible area for both main chain and side chain was calculated with Naccess (Hubbard, S. J. and Thornton, 1993). Our python script requires the Naccess file in the rsa format to automatically remove all the residues with a relative solvent accessible area below 40% for both the side chain and the main chain.

The monomer-monomer docking calculations were carried out with the HADDOCK software (Dominguez et al., 2003). The residues chosen to drive the docking run were given as active residues (directly involved in the interaction) to generate ambiguous interaction restraints (AIRs) with the default upper distance limit of 2 Å. The water-refined models were clustered based on the fraction of common contacts (Rodrigues et al., 2012), FCC = 0.75, and the minimum number of elements in a cluster of 4. For the docking run starting from crystal structures, chain A was used as the input monomer. The number of models generated for each step of the HADDOCK docking procedure were set as follow: 10000 for rigid-body energy minimization, 400 for semi-flexible simulated annealing and 400 for refinement in explicit solvent. The distance violation analysis was performed on the best cluster and the corresponding output written in the ana_dist_viol_all.lis file. In this file we selected all the residues with a violation larger than 3 Å to generate a subset of AIRs to drive a second docking run. Thus, the second docking run was performed using exactly the same conditions as the first one.

We generated 15 models of monomeric *E. coli* L-asparaginase II using the structure of *Wolinella succinogenes* L-asparaginase (Lubkowski et al., 1996) as a template (PDB ID 1WSA, chain A) using Modeller (Eswar et al., 2007). The two proteins have 55% sequence identity. The resulting template-based models featured a very similar backbone conformation, lower than 1 Å from the *E. coli* crystal, but different side chain orientations. Each model was assessed in protein-protein docking using the same AIRs used in the “crystal P 0.25” runs, with all the AIRs (A-C dimer calculation) and after the removal of the ones already satisfied by the A-C dimer (A-D dimer calculation), respectively. The number of models generated for each step were reduced as follow: 1000 for rigid-body energy minimization, 200 for semi-flexible simulated annealing and 200 for refinement in explicit solvent.

All the RMSD values reported in this work were measured on the Cα atoms.

### Solid- and solution-state NMR data

The L-asparaginase II protein [U- ^13^C, ^15^N] was expressed and purified as previously reported (Cerofolini et al., 2019; Giuntini et al., 2017b, 2017a; Ravera et al., 2016), freeze-dried and packed (ca. 20 mg) into a Bruker 3.2 mm zirconia rotor. The material was rehydrated with a solution of 9 mg/mL NaCl in MilliQ H_2_O; the hydration process was monitored through 1D {^1^H}-^13^C cross-polarization SSNMR spectrum and stopped when the resolution of the spectrum did not change any further for successive additions of the solution (Giuntini et al., 2017b, 2017a; Ravera et al., 2016). Silicon plug, (courtesy of Bruker Biospin) placed below the turbine cap, was used to close the rotor and preserve hydration.

SSNMR experiments were recorded on a Bruker AvanceIII spectrometer operating at 800 MHz (19 T, 201.2 MHz ^13^C Larmor frequency) equipped with Bruker 3.2 mm Efree NCH probe-head. All spectra were recorded at 14 kHz MAS frequency and the sample temperature was kept at ≈ 290 K.

Standard ^13^C-^13^C correlation spectra (Dipolar Assisted Rotational Resonance, DARR) with different mixing times (50, 200 and 400 ms) were acquired using the pulse sequences reported in the literature(Takegoshi et al., 2001). Pulses were 2.6 μs for ^1^H, 4 μs for ^13^C; the spectral width was set to 282 ppm; 2048 and 1024 points were acquired in the direct and indirect dimensions, respectively; 128 scans were acquired; the inter-scan delay was set to 1.5 s in all the experiments.

All the spectra were processed with the Bruker TopSpin 3.2 software package and analyzed with the program CARA (Keller, 2007).

The assignment of the carbon resonances of the 2D ^13^C-^13^C DARR spectra of rehydrated freeze-dried ANSII was easily obtained by comparison with the 2D ^13^C-^13^C DARR spectrum collected on the crystalline and PEGylated preparations of L-asparaginase II (Cerofolini et al., 2019; Ravera et al., 2016).

The experimental data used for the Sod1 protein were taken from deposited solution-state 3D ^1^H-^15^N NOESY-HSQC spectrum (Bertini et al., 2009).

Ambiguous assignment lists of the 2D ^13^C-^13^C DARR and 3D ^1^H-^15^N NOESY-HSQC peaks were generated with the program ATNOS/CANDID (Andreas et al., 2016; Guerry and Herrmann, 2012) by setting the chemical-shift–based assignment tolerances to 0.25 ppm and 0.025 ppm, respectively.

## ACKNOWLEDGEMENTS

Financial support was provided by the European Commission (project no. 777536). We thank Prof. Gaetano Montelione for many useful discussions.

## SUPPLEMENTARY INFORMATION

The python script to perform the protocol can be downloaded at the following <u>**LINK**</u>.

## Supplementary figures

**Figure S1.**
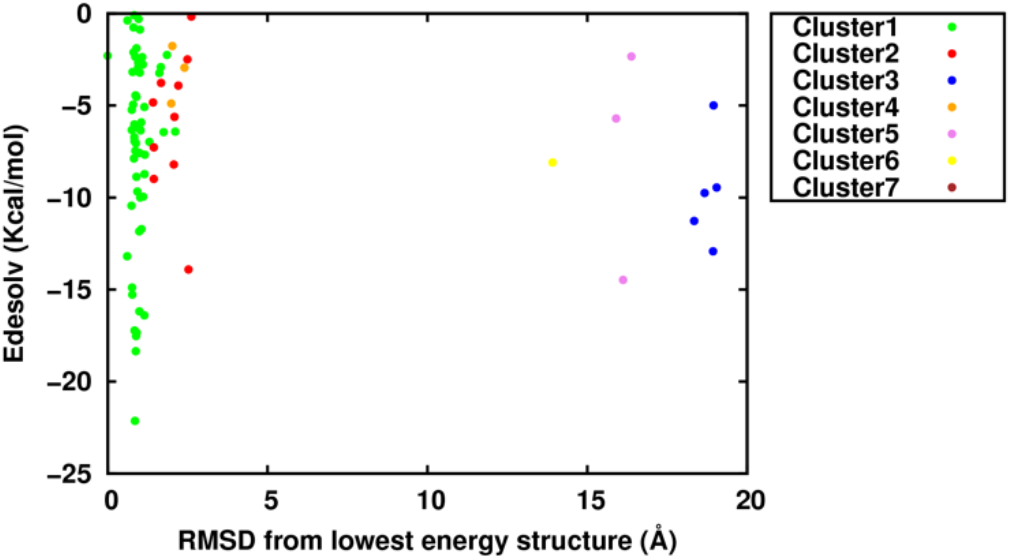
Cluster distribution based on the desolvation energy in the first docking run of L-asparaginase II. The colors of the clusters are the same as in Figure 1.

**Figure S2.**
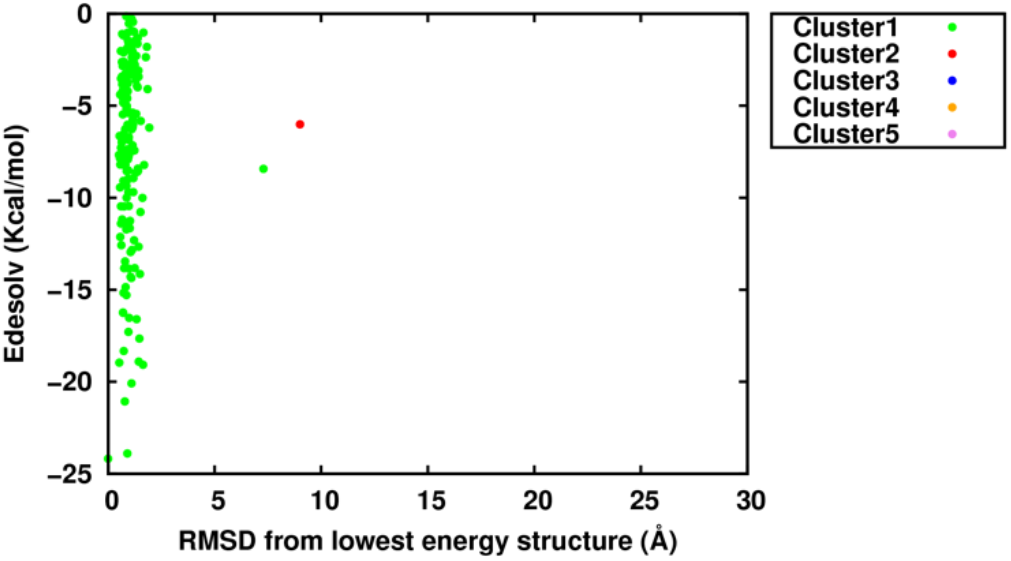
Cluster distribution based on the desolvation energy in the second docking run of L-asparaginase II.

**Figure S3.**
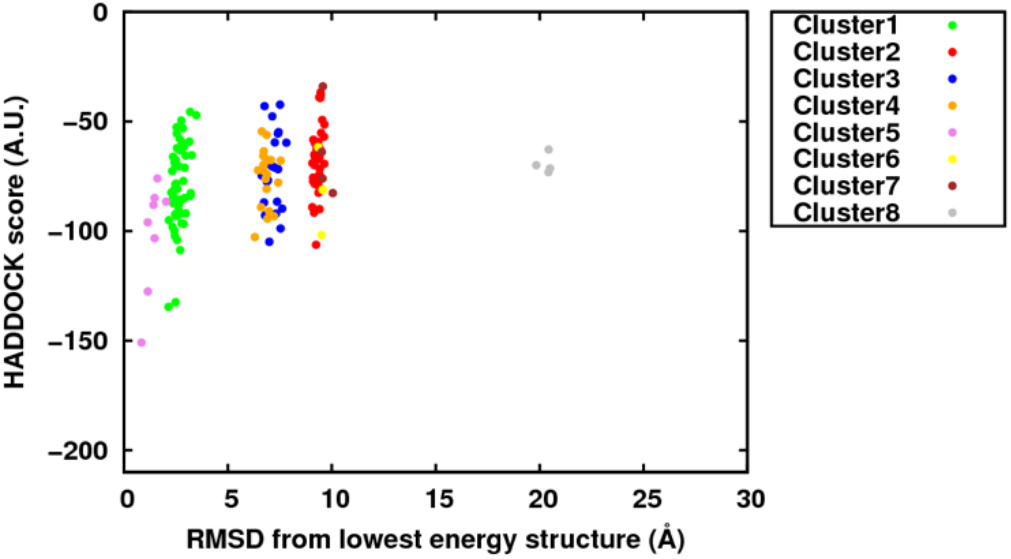
L-asparaginase II clusters distribution obtained from a monomer-monomer docking run performed using the AIRs violated in the third cluster of the first run

**Figure S4.**
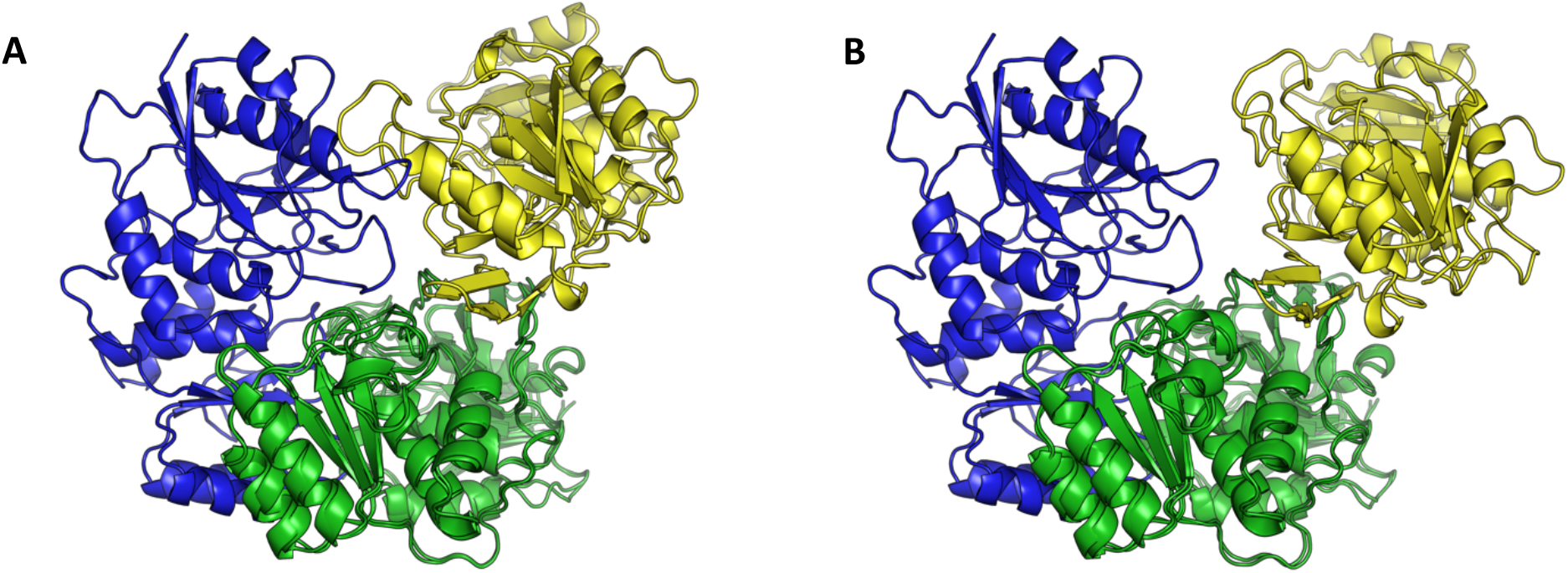
Superimposition on the chain A (in green) of the best L-asparaginase II models. **A**) Model 13 AC dimer is in blue and model 6 AD dimer in yellow. **B**) Mode13 AC dimer is in blue and model 15 AD dimer in yellow.

**Figure S5.**
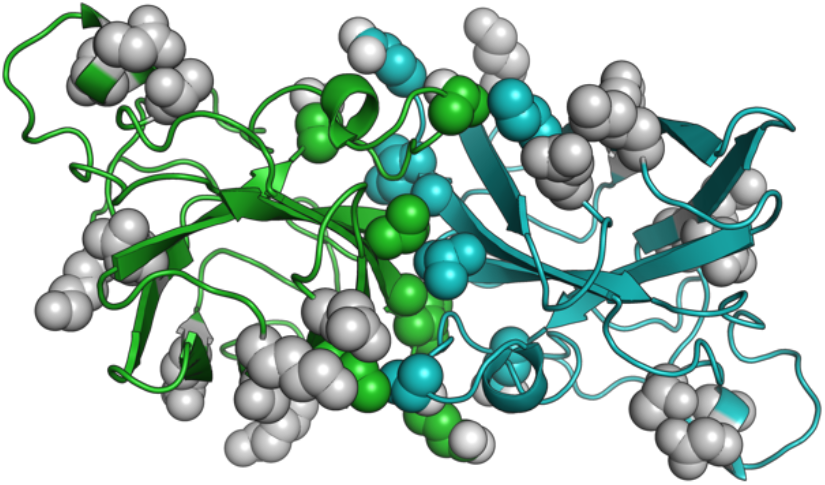
Residues used as AIRS in the docking run of Sod1. Residues forming contacts across the interface are colored as the backbone.

**Figure S6.**
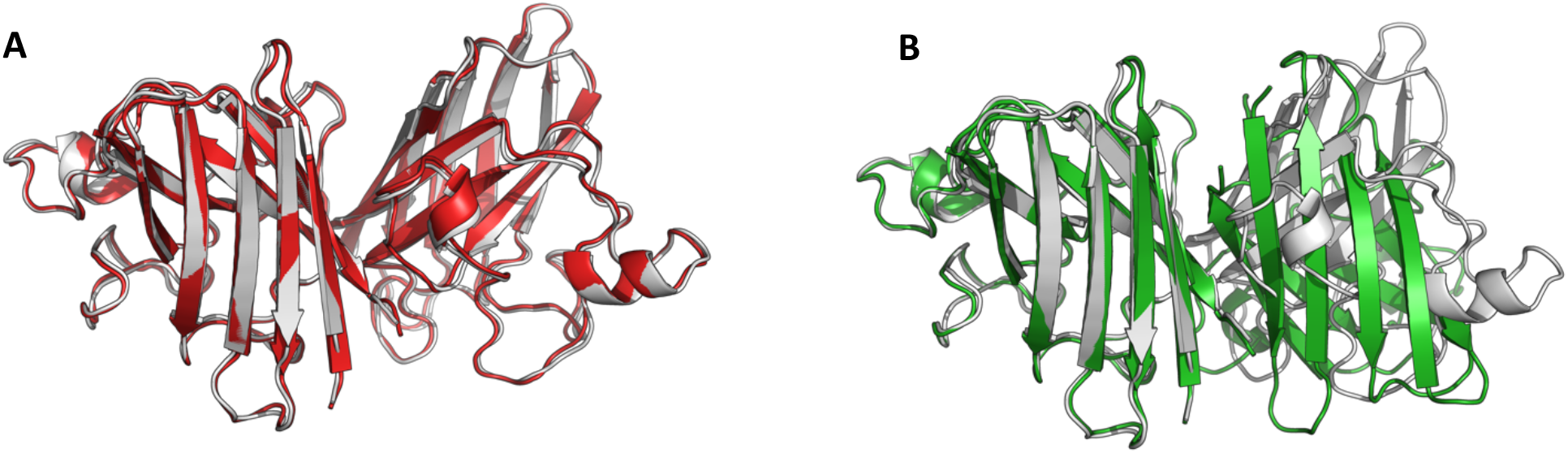
Fitting of the best model of the clusters 1 and 2 on the Sod1 crystal structure. **A**) cluster 2 in red. **B**) cluster 1 in green

